# Whole human organ clearing and multimodal mapping

**DOI:** 10.64898/2026.07.16.738626

**Authors:** Héloïse Policet-Bétend, Tomàs Jordà-Siquier, Geraldine Cuenu, Maeva Badré, Sami Schranz, Julien Da Costa, Céline Brockmann, Pascal Senn, Daniel Huber, Christophe M Lamy

## Abstract

Human anatomy has traditionally been studied using low-resolution approaches such as cadaveric dissection and conventional medical imaging, limiting our understanding of macroscopic structures. Tissue clearing has emerged as a transformative approach, enabling three-dimensional mapping of organs at microscopic resolution, and offering the opportunity to interrogate anatomy across scales. However, its application to human organs remains challenging due to their size, density and optical properties.

Here, we establish multiorgan CleLight (mCleLight) as a generalizable method for clearing, labeling and imaging whole human tissues. mCleLight is broadly applicable across major organ systems and enables the analysis of particularly challenging specimens, including highly heterogeneous samples, dense bones, pigmented tissues and decades-old formalin-fixed archival material. We demonstrate that controlled photoclearing is critical across human tissues both for efficient clearing and for quenching endogenous fluorescence, thereby substantially improving imaging depth as compared to previous methods. mCleLight allows specific labeling and quantitative analysis of complex extended structures such as collagen fibers, vasculature, and innervation, while preserving anatomical relationships, which cannot be easily done with classical histology.

We integrated mCleLight into a pipeline with clinical imaging to achieve a comprehensive, multiscale understanding of organ architecture. We demonstrate the voxel-by-voxel alignment between MRI and microscopy datasets, merging cellular-scale detail and protein-specific labeling with macroscopic medical imaging. By bridging the gap between imaging scales and modalities, mCleLight provides a versatile method for integrated three- dimensional histological mapping and macroscopic anatomy of human organs in health and disease.

## Introduction

Studying the histology and anatomy of human organs is fundamental for understanding their physiology and pathophysiology. Non-invasive imaging techniques such as Computed Tomography (CT), Magnetic Resonance Imaging (MRI) and Positron Emission Tomography (PET) have transformed the study of intact human organs and advanced both clinical diagnosis and biomedical research^1,2^. However, their spatial resolution remains insufficient to resolve the cellular and molecular organization of tissues. Conventional histology overcomes this limitation by providing high-resolution cellular information, but its reliance on thin tissue sections restricts analysis to a two-dimensional plane. This constraint hinders accurate quantification of heterogeneous tissue components, reduces sensitivity for detecting sparsely distributed elements and complicates reconstruction of the three-dimensional organization of extended structures such as collagen bundles, microvascular networks and nerve fibers.

Tissue-clearing techniques have emerged as a powerful strategy to bridge the gap between organ-scale imaging and cellular-resolution histology by enabling three-dimensional visualization of intact tissues. While aqueous- and hydrogel-based clearing methods have been successfully applied to animal models^3,4^, their limited scalability has restricted their application to large human specimens. Organic solvent–based approaches are generally better suited for clearing large tissue volumes^5–8^. However, human tissues present unique challenges for optical clearing, including high autofluorescence, a dense and heterogeneous extracellular matrix, elevated lipid content, pigmentation, calcification, and the effects of prolonged fixation. These factors can compromise tissue transparency, limit antibody penetration, and reduce image quality. Collectively, these challenges highlight the need for a clearing strategy that combines efficient tissue transparency, autofluorescence suppression, deep molecular labeling, and broad compatibility across diverse human tissues.

To address these challenges, we developed mCleLight, a human tissue clearing method that builds upon the CleLight^9^ concept by combining conventional chemical clearing with a photoclearing strategy. Although photobleaching has previously been used to reduce autofluorescence in section-based histology and volumetric human brain^10–15^, CleLight was the first clearing protocol to exploit light exposure not only for autofluorescence quenching but also to enhance tissue transparency. Here, we extend this concept through the incorporation of additional processing steps tissue-specific optimization, enhanced labeling workflows, and systematic validation across a diverse range of human organs.

mCleLight achieves improved tissue transparency, efficient autofluorescence quenching, and deep antibody penetration across a wide range of organ systems, enabling high-resolution volumetric imaging of specimens that are difficult to access using existing approaches, including heavily pigmented, densely calcified, and highly heterogeneous tissues. We demonstrate its broad applicability through the clearing and imaging of intact organs spanning diverse anatomical and structural complexities, including the heart, kidney, finger, inner ear, and ovary. Importantly, mCleLight is compatible with archived clinical specimens, enabling three-dimensional histopathological analysis of retrospectively collected biopsies and autopsy tissues.

Beyond tissue clearing itself, a major challenge is integrating cellular-scale information with anatomical context across larger spatial scales. To address this, mCleLight was integrated into a multimodal imaging pipeline alongside complementary modalities, such as photogrammetry and MRI. This framework enables seamless analysis across spatial scales, linking cellular and molecular features revealed by volumetric microscopy to tissue-, organ-, and whole-body anatomy. By bridging cellular-resolution microscopy with organ-scale imaging, mCleLight provides a framework for comprehensive three-dimensional histology of human organs in both biomedical research and clinical pathology.

## Results

### Photoclearing for multimodal analysis of whole human organs

The mCleLight pipeline was developed to enable a comprehensive, multimodal visualization of whole organ anatomy by integrating tissue clearing with complementary imaging techniques (Fig. 1A). To preserve anatomical information across spatial scales, the pipeline begins with photogrammetric reconstruction of the organs to generate three-dimensional macroscopic surface models, followed by MRI for visualization of internal structures. Finally, chemical and photoclearing enable high-resolution optical imaging for subsequent quantitative analysis.

**Figure 1.**
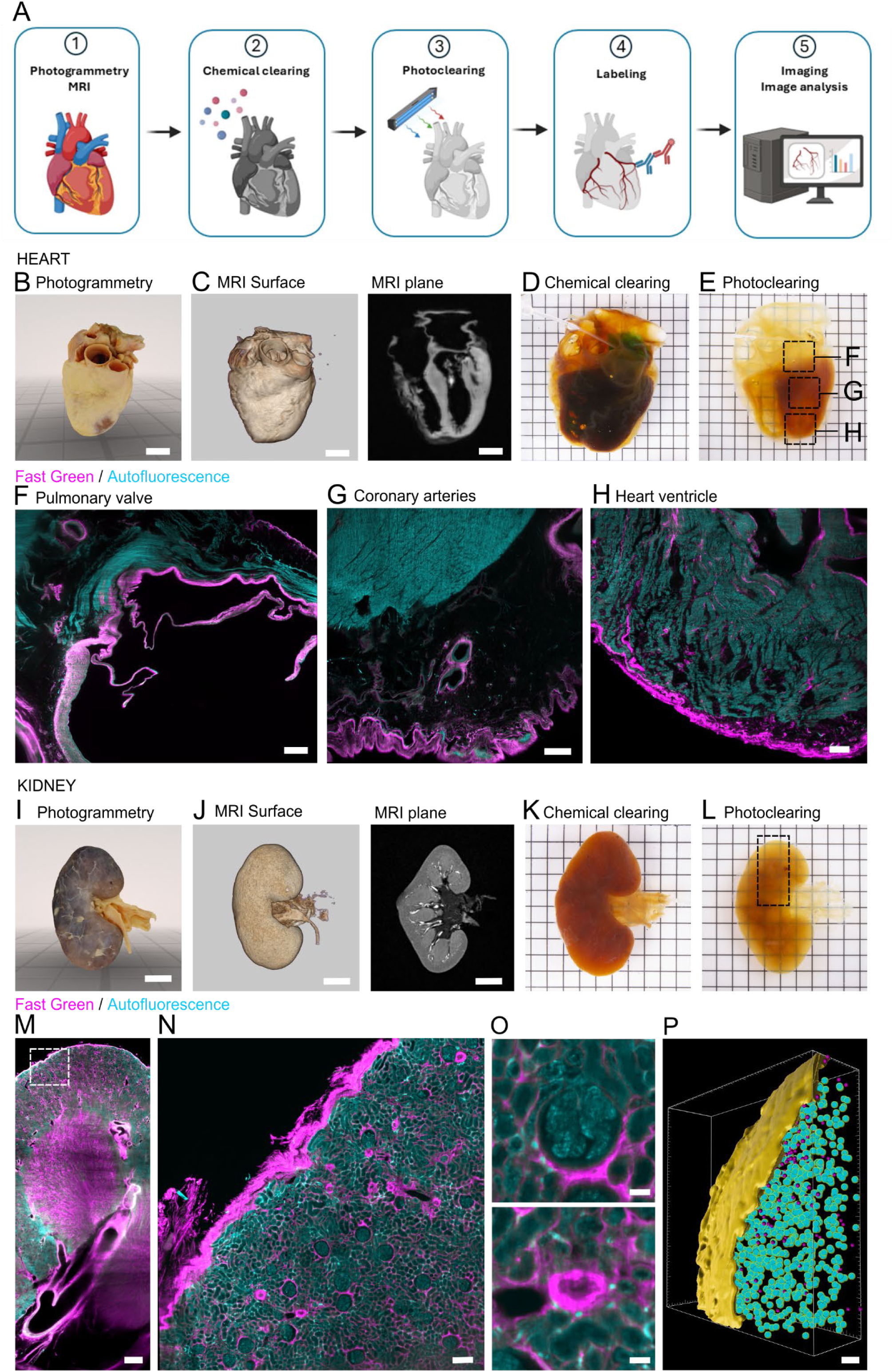
- Photoclearing of whole human organs. **(A)** The mCleLight pipeline involves photogrammetry and clinical imaging to identify anatomical landmarks. The clearing protocol includes two main steps: chemical clearing to delipidate, decolorize, and permeabilize the tissue, followed by photoclearing with a high-power, wide-spectrum LED source to bleach pigments and autofluorescence. Cleared samples are then labeled, imaged using a light-sheet microscope, and processed for quantitative analysis. **(B)** 3D volume reconstruction of a whole human heart with photogrammetry. Scale bars, 2 cm. **(C)** Surface volume rendering of a T1 MRI acquisition of the heart (right panel) and single transversal plane image extracted from the MRI volume (right panel). Scale bars, 2 cm. **(D)** Chemical clearing of the entire heart. A fabric was inserted inside large vessels and cavities to maintain its shape during clearing process. Grid scale 1 cm. **(E)** Heart after mCleLight photoclearing. Grid scale, 1 cm. **(F)** Single plane image from a 3D stack of the inset in panel E showing the pulmonary valve. The entire heart was labeled with Fast Green, while a weak residual autofluorescence signal was imaged to provide additional anatomical contrast. Scale bar 3000 µm. **(G)** Single plane image from a 3D stack of the inset in panel E showing the coronary arteries. Scale bar 500 µm. **(H)** Single plane image from a 3D stack of the inset in panel E. The ventricular myocardium was resolved. Scale bar 500 µm. **(I)** 3D volume reconstruction of a whole human kidney with photogrammetry. Scale bars, 2 cm. **(J)** Surface volume rendering of a T2 MRI acquisition of the kidney (left panel) and single transversal plane image extracted from the MRI volume (right panel). Scale bars, 2 cm. **(K)** Chemical clearing of the entire kidney. Grid scale 1 cm. **(L)** Kidney after mCleLight photoclearing. Grid scale, 1 cm. **(M)** The entire kidney was labeled with Fast Green and was imaged after mCleLight. Background autofluorescence signal was imaged to provide additional anatomical contrast. Scale bar, 2000 µm. **(N)** Higher magnification of the inset in panel K. Degenerated glomeruli were visualized by their spherical shape and fibrous composition labeled with Fast Green. Scale bar, 300 µm. **(O)** Single plane images form 3D stacks of a non-degenerated glomerulus (upper panel) and a degenerated glomerulus (lower panel). Scale bars 20 µm. **(P)** 3D volume segmentation in the renal cortex showing 20% of degenerated and 80% non-degenerated glomeruli. Scale bar, 500 µm.

We first applied the pipeline to a whole human heart. Following dissection, photogrammetry modeled the major vessels (Fig. 1B), while MRI revealed cardiac chambers and wall thickness (Fig. 1C, Supplementary Fig. 1A). Because clearing whole human organs is particularly challenging due to their structural heterogeneity, density, and pigmentation, we adapted the CleLight protocol^9^ by extending dehydration and delipidation times in methanol and dichloromethane, as well as the quadrol ammonium treatment step for effective decolorization. In addition, the heart was cleared using a perfusion-based approach to improve reagent penetration throughout the tissue (Supplementary Fig. 1B-C).

Solvent-based chemical clearing alone provided limited transparency (Fig. 1D). Therefore, we introduced photoclearing to enhance tissue transparency using a high-power, wide-spectrum LED source, providing illumination approximately three times brighter than direct sunlight. Photoclearing resulted in marked depigmentation and substantially improved transparency, even with intrinsic tissue pigmentation from blood or muscle fibers (Fig. 1E; Extended data 1). Whole-heart photoclearing was followed by Fast Green staining, enabling microscopic visualization of collagen distribution. We visualized prominent collagen localization within the fibrosa layer of the pulmonary valve (Fig. 1F). Intense collagen staining was also observed in the pericardium. The coronary arteries gave rise to perforating branches that enter the myocardium and coursed between the cardiac muscle fibers (Fig. 1G-H).

To demonstrate the versatility of the pipeline, we next applied mCleLight to a whole human kidney. Photogrammetry reconstructed the renal hilum, clearly delineating the renal artery, renal vein and ureter, while MRI revealed the cortex, medullary pyramids, interlobar and interlobular vessels (Fig. 1I-J). Following tissue clearing, high-resolution imaging of collagen structures revealed fibrotic, degenerated glomeruli (Fig. 1K-O; Extended data 2). Residual autofluorescence was used to identify non-degenerated glomeruli, enabling volumetric segmentation of glomerular degeneration (Fig. 1P). About 20% of glomeruli were degenerated.

Overall, the introduction of photoclearing is a key feature of mCleLight to enhance transparency in cleared organs. Together with photogrammetry and MRI, mCleLight provides a versatile multimodal framework for three-dimensional anatomical imaging of whole human organs.

### Autofluorescence quenching of human organs

Human organs are structurally dense and contain diverse pigments, as well as age-dependent autofluorescent molecules, making clearing and imaging particularly challenging. We therefore sought to determine whether photoclearing could substantially reduce intrinsic autofluorescence across large human organs in addition to improving tissue transparency.

To assess this, we selected human specimens with diverse compositions, sizes, and pigmentation challenges. These included a 1 cm-thick lung section containing anthracotic pigments, 1 cm-thick liver and kidney sections rich in hemoglobin and heme-derived pigments, highly collagenous carotid artery segment, half-thyroid specimen, and whole eye containing melanin-rich structures (Fig. 2; Supplementary Fig. 2). We quantified autofluorescence in unstained chemically cleared samples before and after high power broadband illumination. The photoclearing process induced substantial photobleaching, resulting in a marked reduction of autofluorescence intensity across all tested excitation wavelengths. This reduction was consistently observed across human specimens spanning a broad range of sizes, compositions, and pigment content.

**Figure 2.**
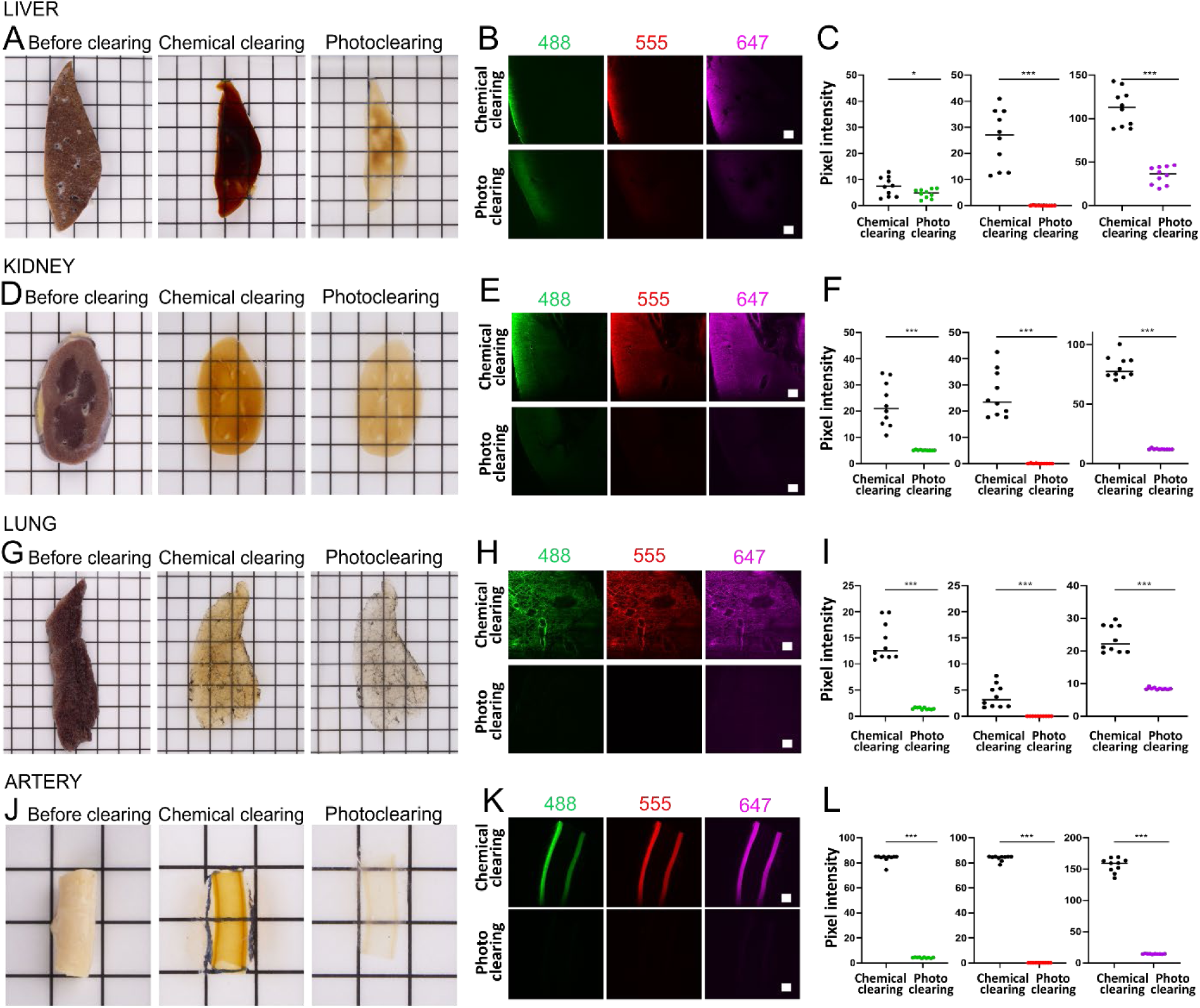
– Autofluorescence quenching across human organs. **(A)** Liver 1cm-thick section before clearing, after chemical clearing and after photoclearing. Grid scale 1 cm. **(B)** Imaging of autofluorescence levels after chemical clearing (upper rows) and after photoclearing (lower rows) across different excitation wavelengths (488, 555 and 647nm). The same imaging and image rendering parameters were used after chemical clearing and after photoclearing. Autofluorescence levels were significantly reduced. Scale bars, 500 µm. **(C)** Quantification of autofluorescence levels after chemical clearing and photoclearing. Values represent average pixel intensity in arbitrary units (A.U.) over ten orthogonal views through the width of the organs. **(D)** Kidney 1cm-thick section before clearing, after chemical clearing and after photoclearing. Grid scale 1 cm. **(E)** Imaging of autofluorescence levels after chemical clearing (upper rows) and photo clearing (lower rows). Scale bars, 500 µm. **(F)** Quantification of autofluorescence levels after chemical clearing and after photo clearing. Values represent average pixel intensity in arbitrary units (A.U.) over ten orthogonal views through the width of the organs. **(G)** Lung 1cm-thick section before clearing, after chemical clearing and after photoclearing. Grid scale 1cm. **(H)** Imaging of autofluorescence levels after chemical clearing (upper rows) and after photo clearing (lower rows). Scale bars, 500 µm. **(I)** Quantifications of autofluorescence levels after chemical clearing and after photo clearing. Average pixel intensity (A.U.). **(J)** Common carotid artery segment before clearing, after chemical clearing and after photoclearing. Grid scale 1 cm. **(K)** Imaging of autofluorescence levels after chemical clearing (upper rows) and after photo clearing (lower rows). Scale bars, 500 µm. **(L)** Quantification of autofluorescence levels after chemical clearing and photo clearing. Values represent average pixel intensity in arbitrary units (A.U.) over ten orthogonal views through the width of the organs.

Together, these results demonstrate that photoclearing enables superior optical clarity in large, highly fibrous, and structurally complex organs while reducing autofluorescence across all wavelengths and improving light-sheet penetration depth.

### Deep dye staining of human organs

Deep dye staining is crucial for investigating the cellular and molecular architecture of large organs. To demonstrate mCleLight’s versatility, we assessed its ability to stain a range of human organs and reveal their spatial organization.

We explored highly vascularized organs by clearing a full archival thyroid lobe. Using the nuclear dye TO-PRO-3 combined with lectin labeling, we resolved the 3D vascular network organized around colloid-filled thyroid follicles (Fig. 3A-C). Clearing and staining of an archival esophagus slice revealed blood vessels arranged around tubulo-alveolar mucous glands within the submucosa (Fig. 3D-F). Similarly, in the pancreas, staining with the small-molecule dyes eosin and Fast Green delineated the 3D organization in lobules separated by septa containing arteries (Supplementary Fig. 3A).

**Figure 3.**
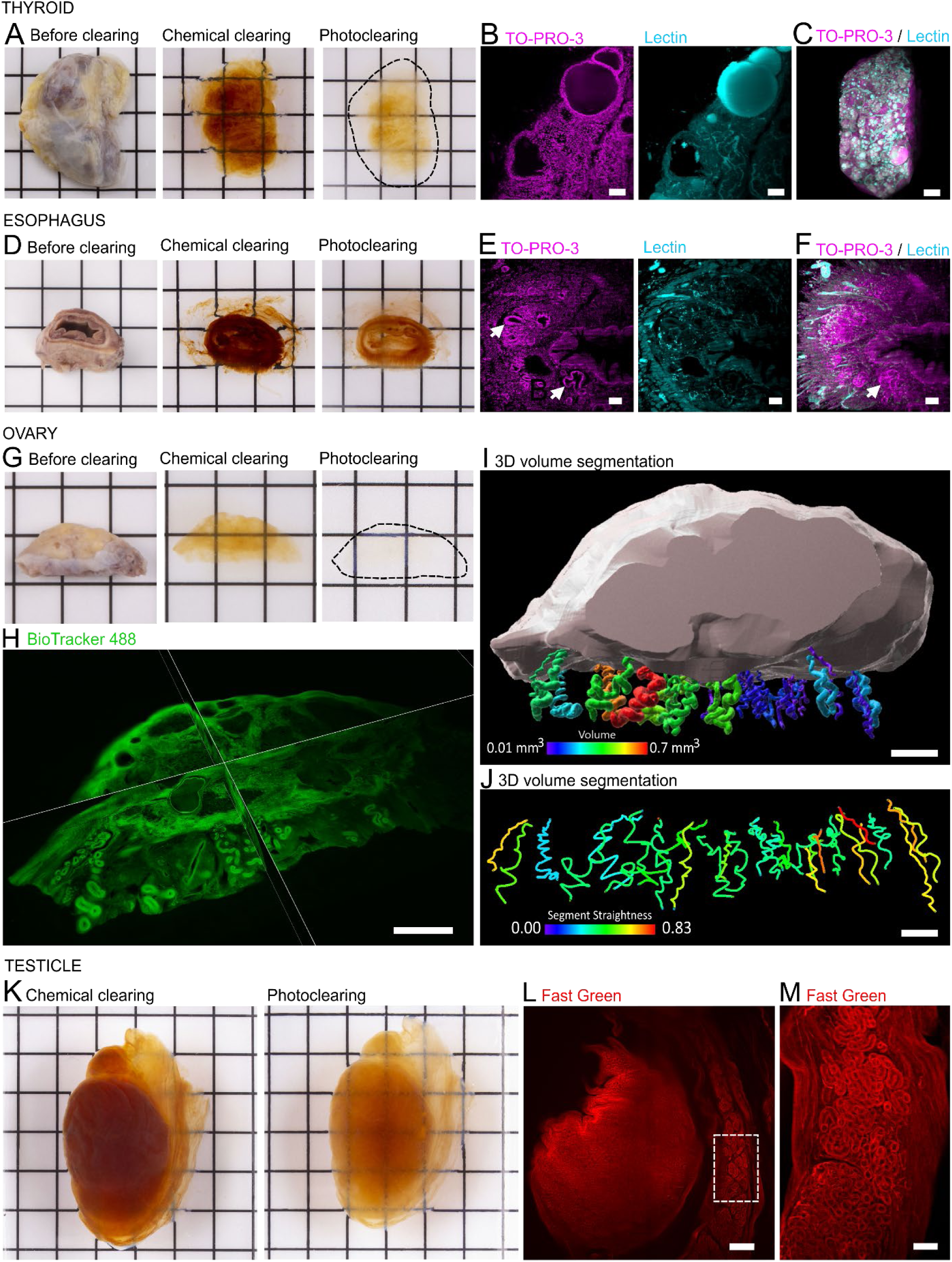
- Deep staining of human organs. **(A)** Archival thyroid gland lobe before clearing, after chemical clearing and after photoclearing. Grid scale 1 cm. **(B)** Maximum intensity projection, planes extracted from a 3D stack of an archival thyroid sample stained with TO-PRO-3 and lectin. Scale bar, 100 µm. **(C)** 3D volume reconstruction of the archival thyroid sample with merged labeling for TO-PRO-3 and Lectin. Scale bar, 500 µm. **(D)** Archival esophagus 1cm-thick section before clearing, after chemical clearing and after photoclearing. Grid scale 1 cm. **(E)** Maximum intensity projection from a 3D stack subset and **(F)** 3D volume reconstruction of an archival esophagus tissue stained with TO-PRO-3 and lectin. Esophageal glands are indicated by arrows. Scale bar, 100 µm. **(G)** Entire ovary before clearing, after chemical clearing and after photoclearing. Grid scale 1 cm. **(H)** Sagittal and frontal planes allow the visualization of the inner structure of the ovary. Scale bar, 2 mm. **(I)** 3D volume segmentation of the ovary and its blood vessels at the level of mesovarium. Color scale volume: purple=0.1mm3 to red=0.7 mm3. Scale bar, 1.5 mm. **(J)** 3D Quantification of vessels’ straightness in the ovary. Scale bar, 1 mm. **(K)** Entire testicle before clearing, after chemical clearing and after photoclearing. Grid scale 1 cm. **(L)** Maximum intensity projection from a 3D stack subset of a testicle and epididymis labelled with collagen staining. Scale bar, 2 mm. **(M)** Inset showing the body of the epididymis and its tubules at higher magnification. Scale bar, 200 µm.

In the female reproductive system, we cleared and stained an entire ovary using nuclear marker BioTracker 488 (Fig. 3G). In a sagittal and frontal 3D reconstruction, we could visualize the inner structure of the ovary, identifying postmenopausal follicles and convoluted vessels entering the hilum (Fig. 3H). Large vessels displayed an intricate architecture, which was reconstructed in 3D and segmented to determine their volume, size and straightness (Fig. 3I-J; Extended data 3). In the male reproductive system, we cleared and labeled collagen structures in the testis (Fig. 3K). We identified complex intricate network of seminiferous tubules in the testis and epididymis (Fig. 3L-M).

We further co-stained a lung section using eosin, which labels basic structures, and Fast Green, which stains collagen. We revealed the 3D architecture of terminal and lobular bronchioles, as well as the extracellular matrix of the alveolar septa and sacs (Supplementary Fig. 3B-C). Collagen-rich pleural surface and interlobular septa were particularly prominent (Supplementary Fig. 3D, Extended data 4). In the kidney, we used a lectin dye as a blood vessels marker, which delineated the vasa recta and the interlobar and interlobular vessels (Supplementary Fig. 2E-F). Segmentation analysis showed that vessels occupied 58% of cortical volume, whereas the vasa recta accounted for 41% of the outer stripe of the medulla (Supplementary Fig. 3G-H). In the liver, collagen labeling enabled identification of large pathological fibrotic regions (Supplementary Fig. 3I-J).

Together, these results show that mCleLight supports deep staining of heterogeneous organs, enabling 3D segmentation and quantitative analysis of tissue architecture while revealing clinically relevant structures.

### Deep immunolabeling for mapping of human organs

While small-molecule dyes reveal overall tissue and vascular architecture, they are inadequate for identifying specific protein expression patterns. The next step was to perform immunostaining, which requires efficient tissue permeabilization to achieve uniform antibody penetration throughout intact organs. Previously, we demonstrated that chemical treatments based on guanidine hydrochlorid and acetic acid, which soften the extracellular matrix, lead to homogeneous staining of large tissue samples^9^. We applied these treatments to whole organs; however, acetic acid treatment was omitted for highly fibrous or cavity-containing tissues, such as the eye and the lung as it induced increased shrinkage and collapse of internal structures.

Using this adapted permeabilization strategy, we assessed the feasibility of deep immunolabeling in whole human organs. In the kidney, aquaporin 2 immunolabeling enabled the visualization of the renal tubules, which traveled parallel to the interlobular arteries (Fig. 4A). Glomeruli were also visualized with nuclei stain TO-PRO-3 in between renal tubules (Fig. 4B). In the lung, smooth muscle actin (SMA) immunolabeling enabled 3D visualization of airways and arteries (Fig 4C-D).

**Figure 4.**
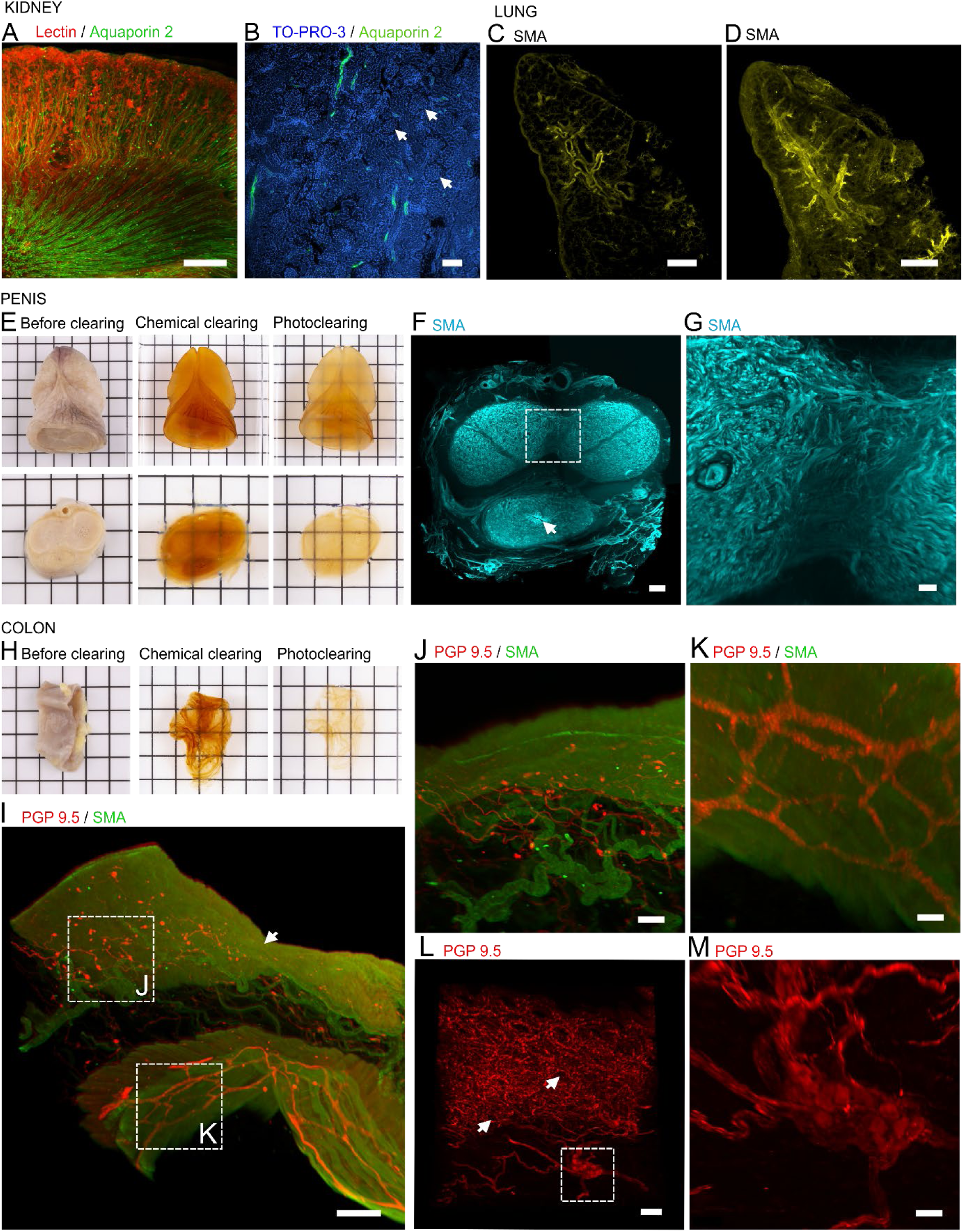
– Deep immunolabeling of human organs. **(A)** 3D volume reconstruction of a kidney section with nephron tubules immunolabeled with aquaporin 2 and vessels labeled with lectin. Scale bar, 1.5 mm. **(B)** Single plane image extracted from a 3D stack of a kidney section labeled for nephron tubules marker Aquaporin 2 and nuclear stain TO-PRO-3. Glomeruli could be identified with the nuclear staining (arrow). Scale bar, 100 µm. **(C)** Single plane image extracted from a 3D stack of SMA immunolabeling in the lung. Scale bar, 1 mm. **(D)** 3D volume reconstruction of SMA labeling in the lung. Scale bar, 1 mm. **(E)** Penis glans (upper panel) and penis 1-cm thick coronal section (lower panel) before clearing, after chemical clearing and after photoclearing. Grid scale 1 cm. **(F)** 3D volume reconstruction of a segment of the panel (I) coronal penis section. Tunica albuginea and cavernous bodies are labeled for collagen with Fast Green. Scale bar, 500 µm. **(G)** 3D volume reconstruction of a coronal section of the penis. Blood vessels of the cavernous and spongious bodies were immunolabeled for SMA. The urethra could be visualized (arrow). Scale bar, 2 cm. **(H)** Colon transverse section before clearing, after chemical clearing and after photoclearing. Grid scale 1 cm. **(I)** 3D volume reconstruction of a human colon sample immunolabeled for SMA and PGP 9.5. The side of the mucosae is indicated by an arrow. Scale bar, 800 µm. **(J)** Inset from panel (I) with higher-magnification image of the sub-mucosal plexuses. Scale bars, 100 µm. **(K)** Inset from panel (I) with higher-magnification image of the myenteric plexuses. Scale bars, 400 µm. **(L)** 3D volume reconstruction the mucosal innervation around colonic crypts (arrows). Scale bar, 70 µm. **(M)** Inset showing details of a submucosal plexus. Scale bar, 30 µm.

Novel clearing methods can also facilitate the study of under-examined human anatomical regions. We successfully cleared a whole penis glans (Fig. 4E). We performed immunolabeling for SMA in the coronal slice of a penis to identify smooth muscles, which are essential for the erection and vascular control of the organ (Fig. 4F). We identified vessels connecting both cavernous bodies in 3D (Fig. 4G).

We further cleared a large colon segment (Fig. 4H). While the neuronal protein gene product 9.5 (PGP 9.5) was used to map the enteric nervous system, SMA immunolabeling highlighted the organization of smooth muscle layers (Fig. 4I; Extended data 5). With this method we could address the 3D structure of submucosal and myenteric plexuses (Fig. 4J-K).

Interestingly, nerve fibers extended in the mucosa, forming a highly ramified network around the crypts (Fig. 4L). Three-dimensional segmentation and quantification of nerve fibers across different layers of the colonic wall further showed a higher nerve fiber density in the mucosa than in the circular and longitudinal muscle layers (Supplementary Fig. 4A-B). Quantitative analysis revealed a higher average neuronal density in myenteric plexuses (15 neurons/ganglion) compared to submucosal plexuses (5 neurons/ganglion) (Fig. 4M; Supplementary Fig. 4C). This data indicates that mCleLight enables the quantitative analysis of complex tissue networks, including organ innervation.

These results demonstrate that immunolabeling can be performed homogeneously in large organs and highlight the utility of mCleLight for detailed structural characterization of previously understudied large human organs.

### Multimodal mapping of the inner ear

Calcified structures pose a major challenge for imaging in light-sheet microscopy. Decalcification is therefore a crucial prerequisite for clearing bones and other calcified tissues, such as aged blood vessels. To test our method in this context, we focused on the petrous bone, the densest bone in the human body, which contains the cochlea and vestibular system.

The first step, based on photogrammetry, provided a macroscopic 3D surface reconstruction of the petrous bone, capturing fine external morphological details (Fig. 5A). We then performed T2-MRI, which provided a global view of the cochlea and vestibular system (Fig. 5B). After decalcification with 10% ethylenediaminetetraacetic acid (EDTA) and subsequent mCleLight clearing, the sample was immunolabeled for neuronal marker beta-III tubulin (Fig. 5C-D). Tissue clearing dataset matched both the photogrammetric and microscopic images. Subsequently, we established a voxel-wise registration between the MRI and light-sheet microscopy datasets, enabling precise correlation of structural information across modalities and resolving key compartments such as the scala tympani, scala vestibuli, and cochlear duct (Fig. E-F). Beta-III tubulin immunolabeling revealed fine and deeply embedded nerve structures that were not detectable by MRI alone.

**Figure 5.**
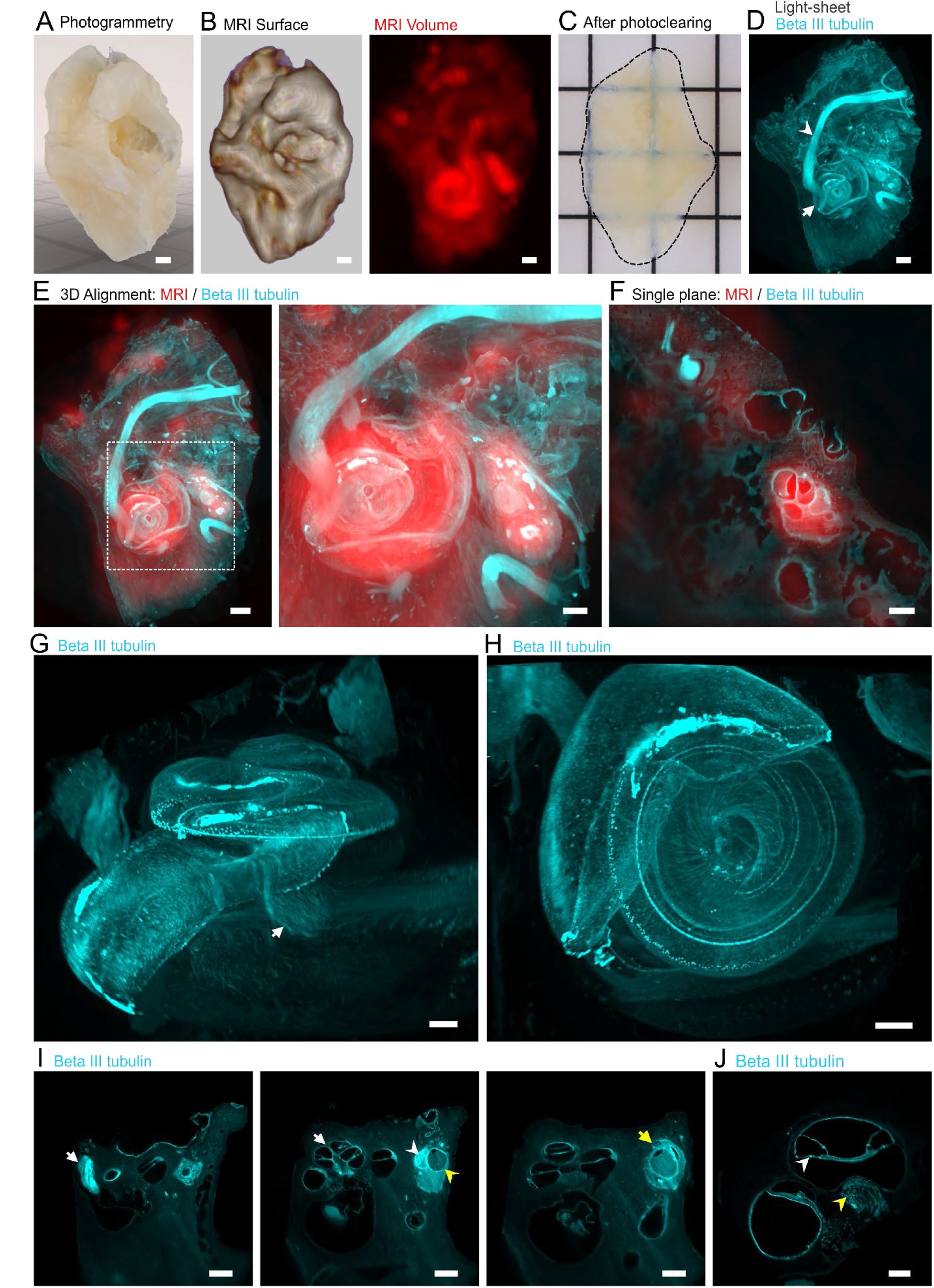
– Multimodal alignment of the inner ear. **(A)** 3D volume reconstruction of the petrous bone with photogrammetry. Scale bar, 2 mm. **(B)** Surface rendering of a MRI T2-acquisition of the petrous bone (left panel) and 3D reconstruction of the same volume (right panel). Scale bars, 2 mm. **(C)** Petrous bone after photoclearing. Grid scale 1cm. **(D)** 3D volume reconstruction of the cleared petrous bone revealing the intrapetrous path of the facial nerve (arrowhead) and the cochlea (arrow). Scale bar, 2 mm. **(E)** Overlaid images of the realigned MRI and light sheet microscopy volumes. Scale bar, 2 mm. Right panel shows a magnified image of the inset indicated on the left panel, Scale bar, 1.5 mm. **(F)** Overlaid single plane image from the realigned MRI and light-sheet microscopy volumes. Scale bar, 1 mm. **(G)** 3D volume reconstructions of the cochlea viewed from the side. The cochlear nerve innervating the cochlea is visible (arrow). Scale bar, 500 µm. **(H)** 3D volume reconstructions of the cochlea viewed from above. Scale bar, 500 µm. **(I)** Single planes images from a 3D acquisition of the cochlea labeled for beta-III tubulin. Left panel shows the facial nerve passing through the petrous bone. Middle panel shows the cochlea (white arrow) divided into the scala vestibuli, media and tympani, together with the utricle (white arrowhead) and a semicircular canal (yellow arrowhead). Right panel shows the ampoullae (yellow arrow). Scale bars, 2 mm. **(J)** Higher-magnification image of the cochlear membranes and cavities, also showing the organ of Corti (white arrowhead) and spiral ganglions (yellow arrowhead). Scale bar, 500 µm.

Tissue clearing three-dimensional reconstruction allowed tracing the intrapetrous portion of the facial nerve and identifying the cochlear nerve (Fig. 5G-H-I; Extended data 6). Tissue clearing enabled a clear visualization of key inner ear structures, including the cochlear compartments of the scala vestibuli, media and tympani, as well as distinct parts of the vestibular system such as the utricle, ampullae and semicircular canals (Fig. 5I). Within the cochlea, the organ of Corti was visible, with immunolabeled afferent fibers of the cochlear nerve and identifiable spiral ganglion cell bodies (Fig. 5J). Combined, tissue clearing and clinical imaging provide a powerful multimodal approach unlocking new opportunities for correlative anatomy and translational studies of complex structures.

### Mapping of musculoskeletal structures

Whole-organ imaging presents a significant challenge due to the intrinsic heterogeneity of biological tissues, which often comprise skin, muscle, bone, cartilage, and connective tissue within a single specimen. To assess the robustness of mCleLight in these conditions, we applied it to an entire human finger and mapped the distribution of Pacinian corpuscles.

We first used MRI to reveal the major anatomical structures, including skin, bone, and flexor tendon (Fig. 6A). Pacinian corpuscles, which are also detectable by MRI^16,17^ were likewise visualized and segmented.

**Figure 6.**
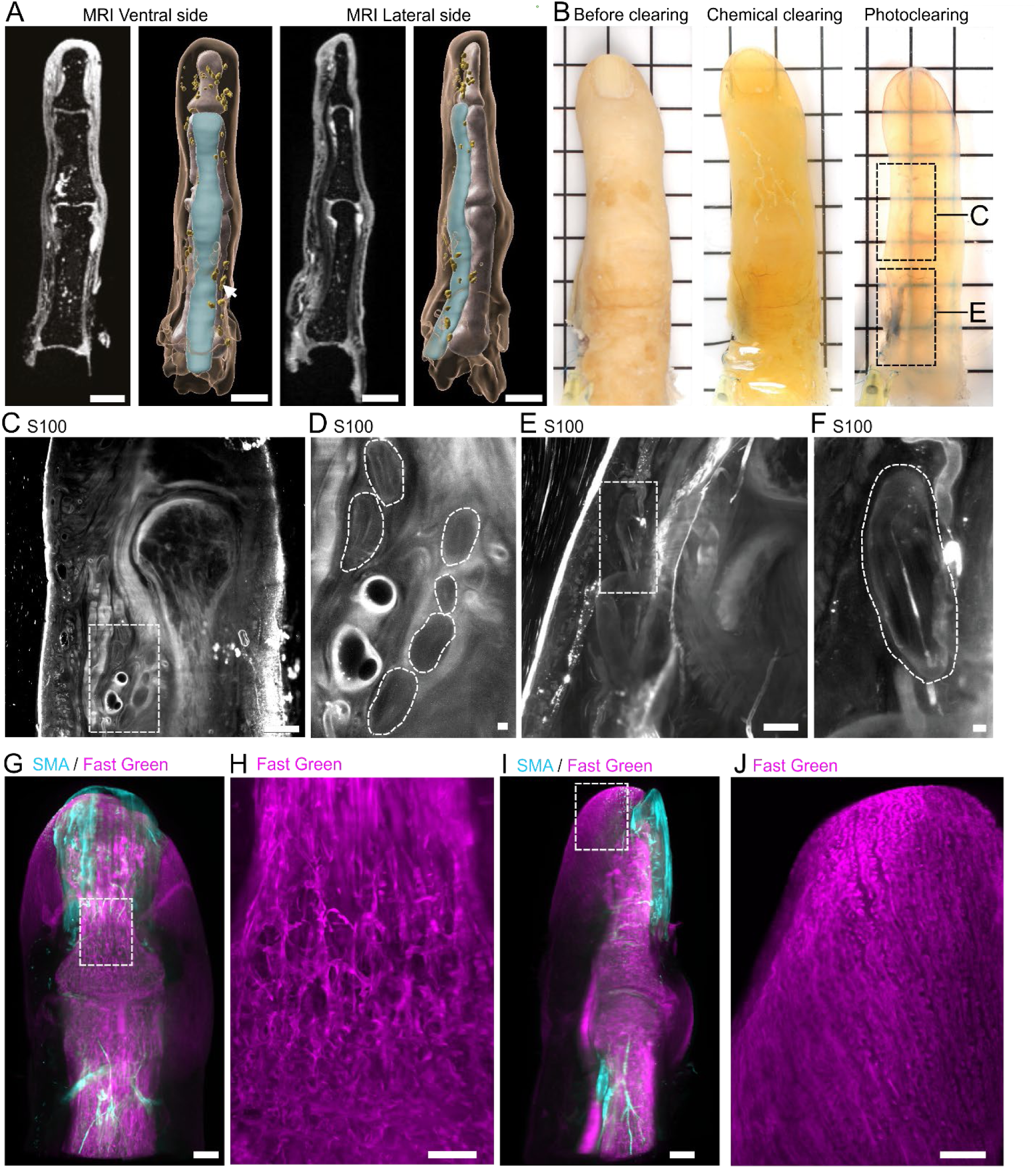
– mCleLight clears musculoskeletal tissues. **(A)** Ventral (left panel) and lateral (right panel) views of a human finger. Left images represent single planes T2-MRI acquisitions and right images show corresponding 3D volume segmentation of the main anatomical structures, including the skin, bone and the flexor tendon (arrow). Scale bar, 1 cm. **(B)** Left ring finger at successive stages of the clearing process: before clearing, after chemical clearing and after photoclearing. Grid scale, 1 cm. **(C)** Longitudinal optical section through the finger from the inset in panel B. Scale bars, 2000 µm. **(D)** Higher-magnification view of individual Pacinian corpuscles of the inset in panel C. Pacinian corpuscles were outlined and observed at various depths, including in the flexor tendon and adjacent to phalangeal bones. The inner core was immunolabeled with S100 antibody. Scale bars, 200 µm. **(E)** Longitudinal optical section through the finger from the inset in panel B. Scale bars, 2000 µm. **(F)** Higher-magnification view of an individual Pacinian corpuscle of the inset in panel E. Pacinian corpuscles were outlined and the inner core was immunolabeled with S100 antibody. Scale bars, 200 µm. **(G)** Dorsal view of a 3D volume reconstruction of a human ring finger with SMA immunolabeling and Fast Green collagen staining. Scale bar, 2 mm. **(H)** Inset showing higher magnification image of the Fast Green collagen staining outlining the trabecular structure of the phalanx. Scale bar, 700 µm. **(I)** Lateral view of a 3D volume reconstruction of a human ring finger with SMA immunolabeling and Fast Green collagen staining. Scale bar, 2 mm. **(J)** Inset showing higher magnification image of the Fast Green collagen staining outlining the fingertip’s reticular collagen network. Scale bars, 1 mm.

Clearing of human fingers began with decalcification, which was verified by computed tomography (CT) showing efficient demineralization of the phalangeal bones (Supplementary Fig. 5A). Subsequent clearing highlighted the critical contribution of photoclearing to the overall clearing outcome (Fig. 6B; Supplementary Fig. 5B-C). The high resolution achieved by light-sheet imaging enabled the visualization and identification of individual Pacinian corpuscles (Fig. 6C–F). S100 immunostaining specifically labeled the inner core. Their characteristic ovoid morphology and surrounding non-fluorescent annular region, corresponding to the fluid-filled spaces between the lamellae of the outer core, further confirmed their identification. Pacinian corpuscles were found along the ventral aspect of the finger close to ligaments and joints.

Beyond mechanoreceptor mapping, we investigated the collagenous and vascular architecture of the finger. Collagen staining delineated the unmineralized bone matrix, corresponding to osteoid matrix. It also revealed the trabecular organization of the distal and middle phalanges, tangential collagen fibers under the skin surface, and radial fibers connecting the skin to the distal phalanx bone (Fig. 6G-J; Extended data 6). SMA labeling revealed the arterial network at the level of the middle phalanx, around the joint and beneath the nail bed. Furthermore, fingertip pulp displayed a dense and highly interconnected collagen reticular network.

Altogether, these findings demonstrate the robustness of mCleLight for clearing complex, bone-containing tissues and its utility for high-resolution mapping of neural, vascular, and connective tissue structures within intact organs.

## Discussion

Resolving the three-dimensional organization of human organs is essential for understanding the structural basis of physiology and disease. Although conventional histology provides cellular resolution, its reliance on thin tissue sections limits the ability to capture the spatial organization of complex tissues over large volumes. Tissue-clearing approaches have transformed volumetric imaging in experimental animal models^18–20^, and increasing efforts have focused on extending these methods to human tissues^21^. However, human specimens remain particularly challenging because of high autofluorescence, heterogeneous extracellular matrix composition, pigmentation, calcification, incomplete antibody penetration and the effects of prolonged fixation. These obstacles have limited the broader adoption of molecularly resolved three-dimensional imaging in human anatomy and pathology.

Here, we present mCleLight, a tissue-clearing protocol that combines conventional chemical clearing with tissue-adapted photoclearing to overcome many of these limitations. Building on our previous demonstration that photoclearing degrades endogenous light-absorbing pigments in post-mortem brain tissue^9^, we show that this principle can be generalized across a broad spectrum of human organs. mCleLight incorporates tissue-specific optimization of photoclearing to accommodate differences in tissue composition, pigmentation and specimen size. These optimizations improve tissue transparency, suppress autofluorescence, enhance imaging depth and enable deep staining and immunolabeling across diverse organ systems. Importantly, mCleLight is a cost-effective protocol which also performs robustly on archival and over-fixed specimens, substantially expanding opportunities for retrospective studies and the analysis of clinically archived material.

Our approach builds upon established tissue clearing strategies, including iDISCO-based and SHANEL-like protocols^6,7,14,22^, which have enabled the clearing and imaging of large biological specimens, including human tissues. However, these methods remain primarily optimized for brain tissue or selected peripheral applications^23–25^, and their performance in whole, structurally complex human organs is often limited by incomplete optical transparency and residual pigmentation. In particular, while previous approaches have demonstrated valuable quantitative analyses in the brain, their extension to intact peripheral human organs has remained challenging. Here, we overcome these limitations by introducing a modified clearing strategy that substantially improves optical transparency in dense human organs. A key advancement is the incorporation of a photoclearing step, which efficiently reduces residual pigmentation and enhances light penetration beyond what is achievable with chemical clearing alone. As a result, our pipeline extends tissue clearing beyond central nervous system applications, enabling robust whole-organ imaging and quantitative analysis across peripheral human tissues.

The versatility of mCleLight expands the range of human tissues accessible to volumetric microscopy. We demonstrate successful clearing and imaging of anatomically challenging specimens, including the heavily mineralized petrous bone, the human finger and reproductive tissues with highly specialized vascular and connective tissue architectures. In particular, volumetric reconstruction of the cochlea and associated nerves provides opportunities to investigate the three-dimensional organization of the human auditory system^17–20^, while imaging of intact finger joints enables detailed characterization of sensory and musculoskeletal structures that are difficult to reconstruct using conventional histology^16^. Similarly, three-dimensional visualization of erectile tissue, such as the corpus cavernosum^31^,offers new possibilities for studying normal and pathological architecture. Collectively, these examples illustrate how mCleLight extends molecularly resolved imaging to human tissues that have remained largely inaccessible using existing whole-mount approaches^32,33^.

Beyond qualitative visualization, mCleLight enables quantitative volumetric analysis of cellular and tissue organization across large human specimens. In the colon, three-dimensional mapping of enteric neurons and nerve fibers over extended tissue volumes provides a framework for studying the spatial organization of the enteric nervous system beyond the limited sampling achievable with conventional histology^34,35^. In the ovary, reconstruction of vascular networks enables quantification of vessel density, diameter and spatial organization, parameters that are directly relevant to angiogenesis during tumor progression^36,37^. More broadly, the ability to extract quantitative structural features from intact tissues creates opportunities for objective characterization of disease-associated remodeling across diverse organ systems.

A key advance of mCleLight is its integration into a multimodal imaging workflow spanning cellular to organ scales. While previous studies have primarily aligned cleared tissues with experimental reference atlases in animal models^38^, integration with human clinical imaging has remained limited^39^. By registering volumetric microscopy datasets to MRI, mCleLight links protein-specific molecular information with macroscopic anatomical context, providing a framework for correlating cellular architecture with clinical imaging features. This multiscale approach has the potential to support the development of comprehensive human organ atlases and facilitate translation between microscopic pathology and non-invasive imaging. Despite these advances, several technical challenges remain. Although improvements in light-sheet microscopy have enabled imaging of increasingly large cleared tissues samples^40^, imaging entire human organs is still constrained by residual light scattering, degradation of light-sheet quality, limited objective working distances and photobleaching. Furthermore, volumetric imaging of human organs generates datasets of unprecedented size, creating substantial demands for data storage, transfer, processing and analysis^41^. Continued developments in imaging instrumentation, computational image analysis and data management will therefore be essential to fully exploit the potential of organ-scale three-dimensional histology.

In conclusion, mCleLight provides a broadly applicable framework for molecularly resolved three-dimensional histology of human organs. By combining tissue-adapted photoclearing, efficient immunolabeling and compatibility with structurally diverse and archival human specimens, the protocol enables comprehensive visualization and quantitative analysis across multiple organ systems. Integration with MRI further extends its utility by connecting cellular-resolution microscopy with organ-scale anatomy, establishing a multimodal platform for investigating human development, physiology and disease. As tissue clearing, imaging technologies and computational analysis continue to advance, mCleLight offers a scalable foundation for building multiscale human organ atlases and expanding the role of volumetric histology in biomedical research and clinical pathology. A bench protocol is available online (https://lamylab.github.io/mCleLight/).

## Materials and methods

### Human tissue samples and dissection

Intact human organs were sourced from the Unit of Anatomy at the University of Geneva from body donors in accordance with the Ethics Committee of the Canton of Geneva (CCER n°2017-01937, CCER). Entire organs were dissected from freshly collected and long-term fixed bodies (Table 1), with careful preservation of nerves and vascularization for subsequent perfusion with PBS 1X Heparin (Braun 2022511). Because coagulated blood exhibits high autofluorescence and opacity, thorough perfusion and blood removal were especially critical in highly vascularized organs such as the penis, heart, and kidney to achieve effective tissue clearing. After washing and blood removal, the organs were fixed by immersion with 4% paraformaldehyde for 48 hours and stored at 4°C in PBS 1X. Except the finger, fixed in a solution of 2% paraformaldehyde and 1% heparin in a close-loop system for 3 days.

**Table 1.** – Specimen data.

| Case | Sex | Age of death (years) | Fixation time | Organs |
| --- | --- | --- | --- | --- |
| Patient 1 | M | 86 | 48h | Kidney, testicle, penis, colon |
| Patient 2 | F | 87 | 48h | Thyroid |
| Patient 3 | F | 91 | 8 years | Esophagus, thyroid |
| Patient 4 | F | 86 | 48h | Lung, esophagus, thyroid, heart, kidney, liver, pancreas, ovary, petrous bone, eye |
| Patient 5 | M | n.k. | 15 years | Penis slice |
| Patient 6 | F | 85 | 48h | Finger |

### Photogrammetry

For the 3D digitization of anatomical specimens, we used a photogrammetric process, a technique that reconstructs three-dimensional digital objects from two-dimensional images by analyzing parallax differences between multiple viewpoints^42–44^. In our study, specimens were suspended on a Silver Serie Mid Turntable (Iconasys), controlled via Shutter Stream software^45^. A total of three sets of 60 pictures were taken (uniform rotation at 6-degree intervals) from an upward oblique angle, frontal view and a downward oblique angle for each model. Image capture was performed using a Nikon Z9 mirrorless camera equipped with a NIKKOR Z 24-70mm f/2.8 S lens, mounted on a tripod. Lighting was provided by two stand-mounted flashes positioned at 45 degrees on either side of the subject, minimizing shadows and enhancing texture details. The entire setup, including the model and turntable, was enclosed in a fabric lightbox to ensure uniform light diffusion and reduce reflections. The captured images were processed using Agisoft Metashape Professional following a standard workflow that included photo alignment, dense point cloud generation, mesh creation, and finally texture mapping.

### Ex-vivo magnetic resonance imaging (MRI)

MRI was performed on the petrous bone, heart, kidney and finger to obtain anatomical images. Prior to imaging, organs were incubated for two days in perfluoropolyether (Galden perfluorinated fluid HT-80, Solvay 31513PP), a proton-free, non-aqueous medium, to prevent motion and avoid susceptibility artifacts during MRI acquisition, except finger immersed in silicone oil. Following incubation, the samples were placed under vacuum conditions to remove trapped air bubbles. Organs were positioned in a transparent container with a custom 3D-printed fixation set (Supplementary Fig. 5C). MRI was performed using a 3T MAGNETOM Prisma Fit scanner (Siemens Healthineers) equipped with a Head/Neck 64-channel coil. All organs were imaged before the clearing process, except for the petrous bone images, which were acquired after clearing. Sequences were optimized for ex-vivo anatomical imaging. Petrous bone images were acquired, using a 3D T2-space transversal sequence (TR: 1860 ms; TE: 93 ms) with an isotropic slice thickness of 0.9 mm. Finger acquisitions were acquired using a 3D T2-space sequence with fat suppression (TR: 1330 ms; TE: 73 ms) with a slice thickness of 0.50 mm. Heart acquisitions were acquired using a 3D T1 DIXCON coronal sequence (TR: 3200 ms; TE: 410 ms) with a slice thickness of 0.94 mm. Kidney acquisitions were acquired using a T2-space sagittal sequence (TR: 3200ms; TE 410ms) with an isotropic slice thickness of 0.49mm. All images were exported in DICOM format.

### Decalcification and CT scan

Highly vascularized organs (heart, kidney, penis) were decalcified 2 weeks in EDTA 10% (NeoFroxx 1108Gr500) pH7 at 37°C by immersion, with renewal of the solution every 3 days (Supplementary Table 2). Bony organs, such as the ear and finger were decalcified by immersion during 3 months in EDTA 20% supplemented with 0.02% sodium azide (Fluka Biochemika 71289) at 37°C, renewed every 7 days. CT imaging was subsequently performed to confirm the absence of residual mineral content using a 64-slice multidetector CT unit (CT VTC; GE Healthcare, Milwaukee, WI, USA). The petrous bone and the finger were scanned using identical acquisition protocols: matrix size 512, field of view 25 cm, slice thickness of 0.625 mm, reconstruction interval of 0.3 mm, tube voltage 100 kVp, tube current 120 mA, rotation time 1 second, and pitch 1.375. Once CT scans showed no remaining bone structures, the tissue-clearing protocol was initiated.

### Perfusion system for heart, kidney, and finger clearing

Heart, kidney and finger were cleared using a closed-loop perfusion system with a peristaltic pump. Right and left coronaries of the heart, as well as the renal artery of the kidney were catheterized and used for perfusion. The fingers were sectioned at the metacarpophalangeal joint and catheterized into the palmar digital artery, secured using three sutures (Ethicon Vicryl 4-0, 1.5 Ph. Eur.) and a drop of medical-grade adhesive. A glass perfusion chamber was sealed with a custom lid and placed on a hot-plate magnetic stirrer. A gap between the holder and the bottom of the chamber allowed placement of a magnetic stir bar, ensuring gentle and uniform agitation of the perfusion medium. Perfusion was driven by a multichannel peristaltic pump, allowing independent perfusion lines. The pump was configured to deliver a continuous, low-pulsation flow approximating the blood flow velocity of 4.8 mL/min reported for human index finger veins^46^.

### Tissue preparation and chemical clearing

Fixed samples were prepared for clearing by PBS 1x washings (P4417-100TAB Sigma Aldrich) with 0.2% Tween-20 (PTw) (PanReac Applichem A4974) and azide 0.02% (Fluka Biochemika 71289) at room temperature (RT).

Eye clearing was performed after removal of the crystalline lens and creation of a corneal incision using a scalpel. Clearing solutions were injected through the incision with a syringe to enhance solution penetration, as the sclera is a dense fibrous tissue that limits passive diffusion. For whole-mouse clearing, the paws were removed prior to clearing to accommodate the dimensions of the light-sheet imaging cuvette. The skin and black fur were preserved throughout the procedure. All incubations of the clearing protocol were done with shaking. The volume of the solutions was at least three times the volume of the organ.

Samples underwent dehydration baths with a gradual increase in methanol (Fisher Chemical M/4000/17) for 1 hour each (50%, 80%, 100%, 100%) at RT. For fatty and dense organs, the first delipidation lasted 2h (50%, 80%) and overnight at 100% and changed as long as the solution was yellow. Samples were rinsed overnight in 66% dichloromethane (DCM) (Fisher Chemical D/1852/17)/ 33% methanol at RT until the sample reached the bottom of the recipient (time is organ-dependent). Samples were washed two times for 30 minutes in 100% methanol and bleached with freshly prepared 5% H_2_O_2_ (Thermo scientific 7722-84-1) in 100% methanol overnight at 4°C. Tissues were rehydrated in serial methanol baths at 80% and 50% for 1 hour each, and a final incubation of PBS 1x Triton (PTx) 0.3% (Sigma-Aldrich T9284) and azide 0.02%.

Subsequently, the tissue was permeabilized for 3 days at 37°C in a solution containing 3% Triton, 0.3M glycine (Pan Reac applichem A1067), 20% (Sigma-Aldrich 41640) and Saponin 10 mg/mL (Sigma-Aldrich S4521) in PBS 1X azide 0.02% refreshed every day.

After two washes in PTw 0.2% azide 0.02%, tissues were incubated for 2 nights at 37°C with a mixture of quadrol at 25% (N,N,N’, N’-Tetrakis(2-hydroxzpropyl)ethylenediamine 98% Sigma-Aldrich 122262-1L) and ammonium hydroxide at 5% (Sigma-Aldrich 30501-1L). Samples were again washed in PTw 0.3% azide 0.02% five times for 30 minutes each at RT.

Subsequently, samples went through sequential methanol baths for 1 hour each, ranging from 50% to 100%, and were left overnight until complete dehydration. Finally, after a 3-hour incubation in 66% DCM/ 33% methanol at RT, and two incubations of 15 minutes at 100% DCM, samples were rendered clear with Dibenzyl ether (DBE) (Sigma-Aldrich 33630) overnight.

### Photoclearing

Samples in DBE were exposed from 2 weeks to 1 month depending on the organ to a white LED light (15000 Im, 4000 K neutral white; Hornbach, Switzerland) at 10°C to avoid overheating (Table 2). The light intensity measured at the sample surface was 350,000 lux, corresponding to an irradiance of 19 mW/cm².

**Table 2.** – Organ-specific decalcification time and light exposure time.

| Organ | Decalcification time | Light exposure time |
| --- | --- | --- |
| Lung | - | 2 weeks |
| Liver | - | 3 weeks |
| Colon | - | 3 weeks |
| Petrous bone | 3 months | 1 week |
| Thyroid | - | 2 weeks |
| Penis | 2 weeks | 3 weeks |
| Ovary | - | 2 weeks |
| Pancreas | - | 2 weeks |
| Eye | - | 3 days |
| Finger (full organ) | 3 months | 2 weeks |
| Heart (full organ) | 2 weeks | 1 month |
| Kidney (full organ) | 2 weeks | 1 month |

### Tissue labeling

Samples were then rehydrated until they reached 50% methanol. After 1 hour in 50% methanol, samples were washed overnight in PTw 0.3% azide 0.02%. At this stage, the extracellular matrix (ECM) of the tissue was loosened by initiating an incubation in 0.5M acetic acid (Merck 1.00244.0500) at RT overnight (except fragile organs like the eye, pancreas) or 3 days incubation for the finger. A second incubation with PBS1x, Triton 3%, 0.05M sodium acetate (Carlo Erba 1310-73-2), 4M Guanidine hydrochlorid 98% (Roth 6069.3) at 37°C overnight for all organs and 3 days incubation for the finger. Samples were washed in PTw 0.3% azide 0.02%.

Samples were blocked overnight at 37°C (5 days incubation for the finger) in PBS 1x, gelatin 0.2% (VWR chemicals 24350.262), 10% DMSO, azide 0.02%. Antibody incubations were done with top up on the 4 first days and incubated for a week further with a primary antibody conjugated with a Fab fragment secondary at 37°C (Table 3). Primary and secondary antibodies were preincubated for 30 minutes at RT before being added to the tissue in a solution of PBS1x-Tween 0.2%, 0.02% azide, 10% CHAPS (Roth 75621-03-3), 1% glycine and 10% DMSO. For the finger, primary and secondary antibodies were incubated separately for 10 days at 37°C in a solution of PBS Tween 0.2%, 3% donkey serum, 5% DMSO, 0.02% azide and 10 mg/L heparin. Chemical dyes were diluted in PBS 1x with a higher NaCl concentration (500mM) and Fast Green was diluted in 100% methanol^47^. After the labeling, the tissues were washed in PTw 0.3% azide 0.02% for one day. Finally, samples were dehydrated in methanol, followed by a 3-hour incubation in 66% DCM/ 33% methanol, and two 15-minute washes in 100% DCM before immersion in DBE. Propyl gallate 0.4% (Sigma Aldrich P3130-100G) can be dissolved in DBE to prevent fluorescence quenching.

**Table 3.** – Antibodies and dyes used with mCleLight.

| Antibody/Dyes | Source | Dilution |
| --- | --- | --- |
| Beta-III tubulin (rabbit) | Abcam ab18207 | 1:100 |
| PGP9.5 (rabbit) | Proteintech 14730-1-AP | 1:250 |
| α-smooth muscle actin (goat) | Abcam ab21027 | 1:500 |
| α-smooth muscle actin (mouse) | Custom made IgG2 clone 1A4, α-SMA, or ACTA<sup>48</sup> | 1:100 - 1:200 |
| Aquaporin 2 (rabbit) | Alomone Lab AQP-002 | 1:500 |
| Androgen receptor (rabbit) | Cell signaling 5153 | 1:300 |
| Donkey anti-mouse FAB fragment | Jackson ImmunoResearch A647 115-607-185 | 1:200 to 1:250 |
| Donkey anti-rabbit FAB fragment | Jackson ImmunoResearch A647 711-607-003 | 1:100 |
| Goat anti-mouse FAB fragment | Jackson ImmunoResearch Cy3 115-167-003 | 1:250 |
| Donkey anti-goat FAB fragment | Jackson ImmunoResearch A647 705-607-003 | 1:500 |
| Lycopersicon Esculentum (Tomato) Lectin (LEL, TL), DyLight 488 | Vector laboratories | 1:100 to 1:250 |
| TO-PRO-3 Iodide | Invitrogen T3605 | 1:100 to 1:1000 |
| BioTracker 488 | Merck millipore SCT120 | 1:100 to 1:500 |
| Fast Green FCF | Thermo scientific A16520.06 | 0.1ng/1mL in 100% methanol<sup>47</sup> |
| Eosin Y | Merck E-4009 | 0.0001% in PBS |

### Light-sheet microscopy imaging

The light-sheet microscope MesoSPIM^49^ features a dual-sided excitation path with a fiber-coupled multiline laser combiner (405, 488, 561, and 647 nm, Toptica MLE). The system includes a 42 Olympus MVX-10 zoom macroscope (0.63X to 3.2X) and a 1× objective (Olympus MVPLAPO 1×).

For imaging, the finger was vertically mounted on a customized holder. Its proximal end at the metacarpophalangeal joint was secured with two screws and iron wire. The mesoSPIM chamber, typically filled with immersion oil, was instead filled with dibenzyl ether to match the refractive index of the cleared tissue. The finger was suspended on the holder and immersed in DBE until most of it was fully submerged. The finger was scanned at 0.8x magnification. To mitigate optical distortions, scattering, and shadowing effects associated with thick and heterogeneous tissue, multi-view imaging was employed to image the finger. Four orientations were used: with the left and right lateral sides facing the detection objective, as well as with the dorsal and ventral sides facing the camera.

Clarity Optimized Light-sheet Microscope (COLM) was custom-designed with two digitally scanned light-sheets with 488, 561, and 647 nm wavelength lasers. Emitted fluorescence was collected with a 10X FLUOR4X / 0.28N.A objective and imaged with an Orca-Flash 4.0 LT digital CMOS camera.

Miltenyi Biotech UltraMicroscope Blaze employed a double illumination light-sheet capable of imaging in 488, 561, and 647 nm laser channels with dynamic focus positioning. We used 1.1X objective (N.A. 0.1), 4X (N.A. 0.35) and 12X (N.A. 0.53) with a filtered 4.2 Megapixel sCMOS camera and a zoom set at 0.66X to 2.75X.

Miltenyi Biotech UltraMicroscope Choros employed a double illumination light-sheet. The system featured a large sample chamber. We used 1.1X objective (N.A. 0.1) and 4X (N.A. 0.35) with a filtered 4.2 Megapixel sCMOS camera.

### Image processing

We used Fiji for single plane image visualization and maximum intensity projections. Imaris (v10 Oxford Instruments) was used for 3D image reconstructions. Fiji plugin Enhance Local Contrast (CLAHE)^50^ was applied on the lung single plane (Supplementary Fig. 3B); esophagus (Fig 3E); Ovary (Fig 3H); Testicle (Fig 3L, M); Kidney (Fig. 4B). All videos were performed with Imaris (v10).

Deconvolution was performed on heart and kidney acquisitions using Huygens Essential (Scientifique Volume Imaging) (Fig 1F-H; 1N-O). Large organs required mosaïc acquisitions, which were stitched according to the microscope and data size acquired: Bigstitcher, Huygens Professional (Scientific Volume Imaging), Imaris Stitcher (Oxford Instruments) or Terastitcher 1.10.18. Finger multi-view acquisitions were manually stitched on Imaris (v10 Oxford Instruments). Clearing and MRI data 3D volumes were manually aligned on Imaris (v10 Oxford Instruments).

The three-dimensional segmentation and quantification of blood vessels and degenerated glomeruli in the kidney, as well as nerve fibers in the colon were performed on raw data using Imaris (v10 Oxfort Instruments) intensity thresholding. Neuronal somata in the colon’s ganglions and non-degenerated glomeruli in the kidney were counted manually. Segmentation of blood in the ovary was done using the Segmentation Editor module of Amira (Thermo Fisher Scientific). Separate label fields were created for each vessel. 3D surface segmentation was exported from Amira to Imaris to extract volumetric information (volume and segment straightness).

### Autofluorescence quantification

Fiji (ImageJ) was used to measure the autofluorescence in samples imaged with a single light-sheet illumination side. Pixel intensity was quantified in ten randomly selected planes throughout the sample, close to the illumination side of the light-sheet. The average pixel intensity across the ten planes was then calculated and compared across different conditions.

### Quantification and statistical analysis

Statistical analyses were performed using GraphPad Prism 10.3. The normality of data distribution was evaluated with the D’Agostino and Pearson tests. For normally distributed data, comparisons were made using an unpaired t-test. Results were deemed statistically significant when P < 0.05.

## Acknowledgments

The authors gratefully acknowledge Marie-Luce Bochaton-Piallat (Faculty of Medicine, University of Geneva) for kindly providing the SMA antibody. The authors also thank Nicolas Liaudet (Bioimaging Core Facility, Faculty of Medicine, University of Geneva) for his expertise in image analysis; Priscilla Soulié (Faculty of Medicine, University of Geneva) for her expertise in kidney histological structures; and Francis Rousset (Faculty of Medicine, University of Geneva) for his expertise in ear anatomy. We acknowledge the Advanced Lightsheet Imaging Center (ALICe) at the Wyss Center for Bio and Neuro Engineering (Geneva). Authors also thank Nicolaas Van Der Voort (Scientific Volume Imaging) for his assistance with deconvolution of large datasets. Finally, the authors sincerely thank Raphael Kurtz and Christine Ahlert (Miltenyi Biotec) for providing the opportunity to perform whole-kidney and whole-heart imaging on the UltraMicroscope Choros and for their expertise throughout the imaging process. Illustrations were created with BioRender.com.

Authors acknowledge fundings from the Swiss National Science Foundation to Christophe Lamy (grant 219457) and Daniel Huber (grant 184829).

