## Supplementary information for "Whole human organ clearing and multimodal mapping"

**Supplementary figures**

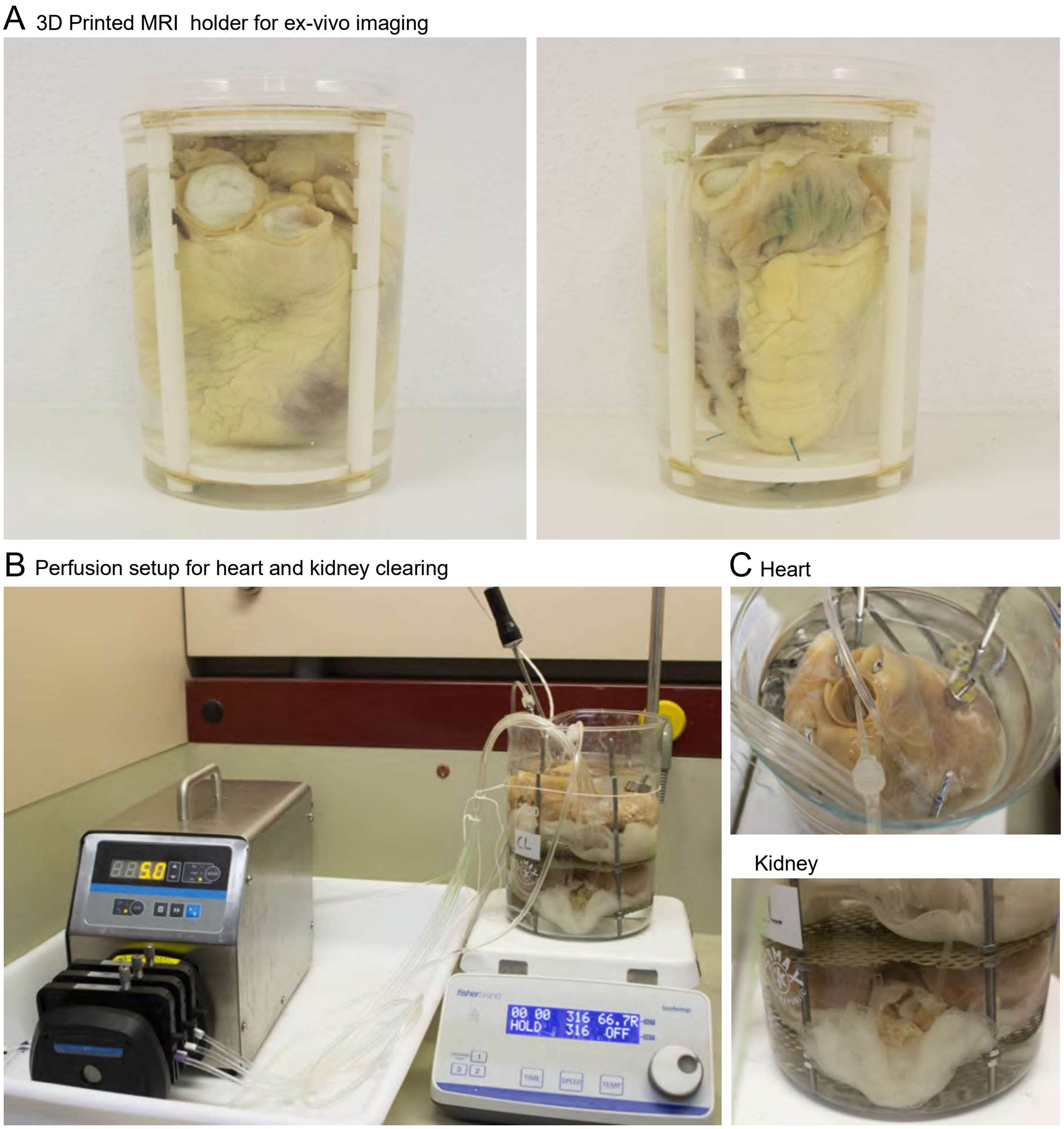


**Supplementary Figure 1 – Processing of whole human organs. (A)** 3D printed holder for ex-vivo MRI imaging of the heart. **(B)** Perfusion setup to clear the whole heart and kidney using a peristaltic pump. **(C)** Details of the perfusion system with catheters in the right and left coronary arteries of the heart and in the renal artery of the kidney.

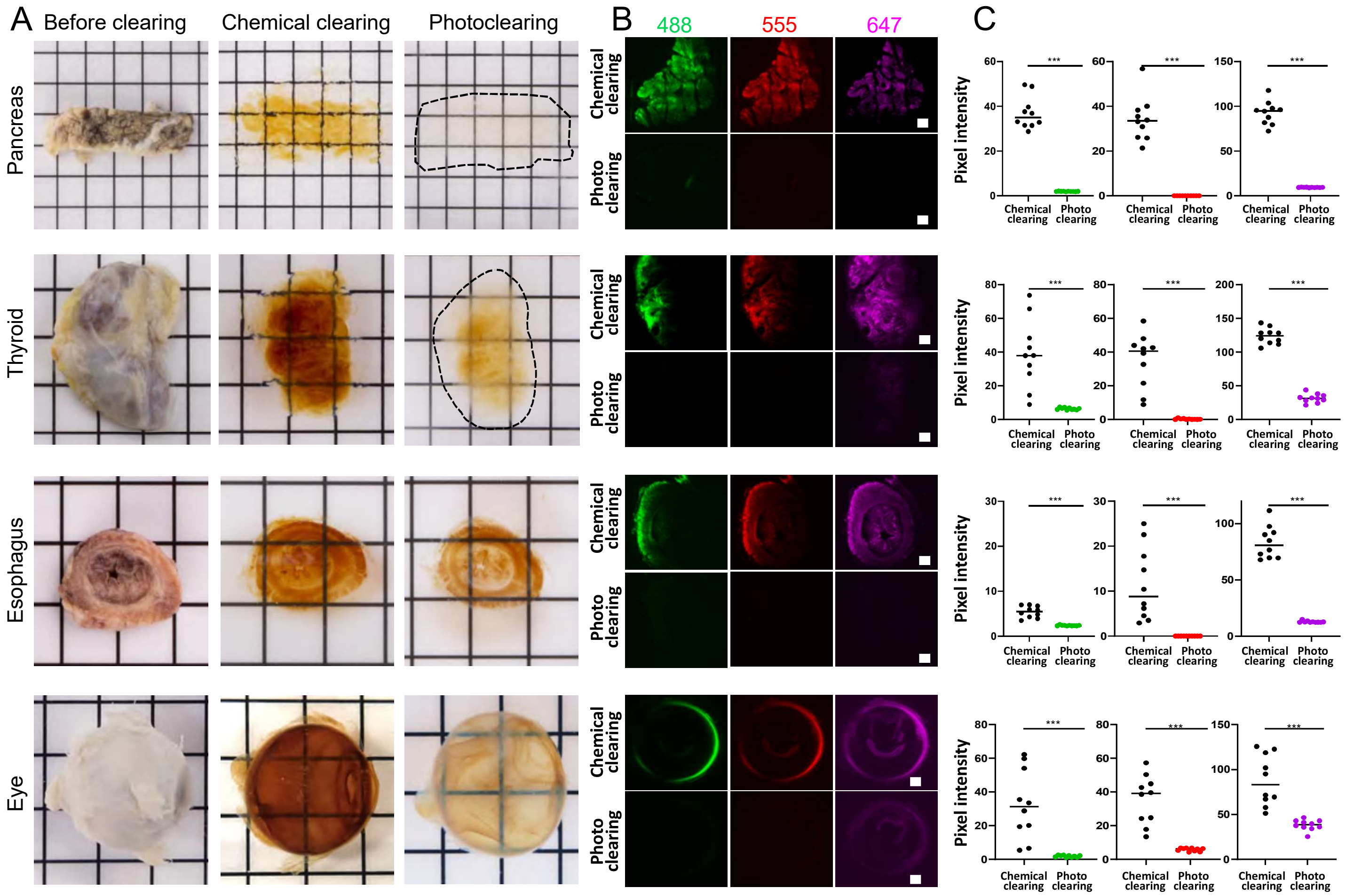


**Supplementary Figure 2 – Effect of photoclearing on tissue transparency and autofluorescence quenching. (A)** Pancreas section (1cm-thick), thyroid lobe, esophagus section (1cm-thick) and eyeball before clearing, after chemical clearing and after photoclearing. **(B)** Imaging of autofluorescence levels after chemical clearing (upper rows) and after photoclearing (lower rows) across different excitation wavelengths (488, 555 and 647nm). The same imaging and image rendering parameters were used after chemical clearing and after photoclearing. Autofluorescence levels were significantly reduced. Scale bars, 500 μm. **(C)** Quantification of autofluorescence levels after chemical clearing and photoclearing. Values represent average pixel intensity in arbitrary units (A.U.) over ten orthogonal views through the width of the organs.

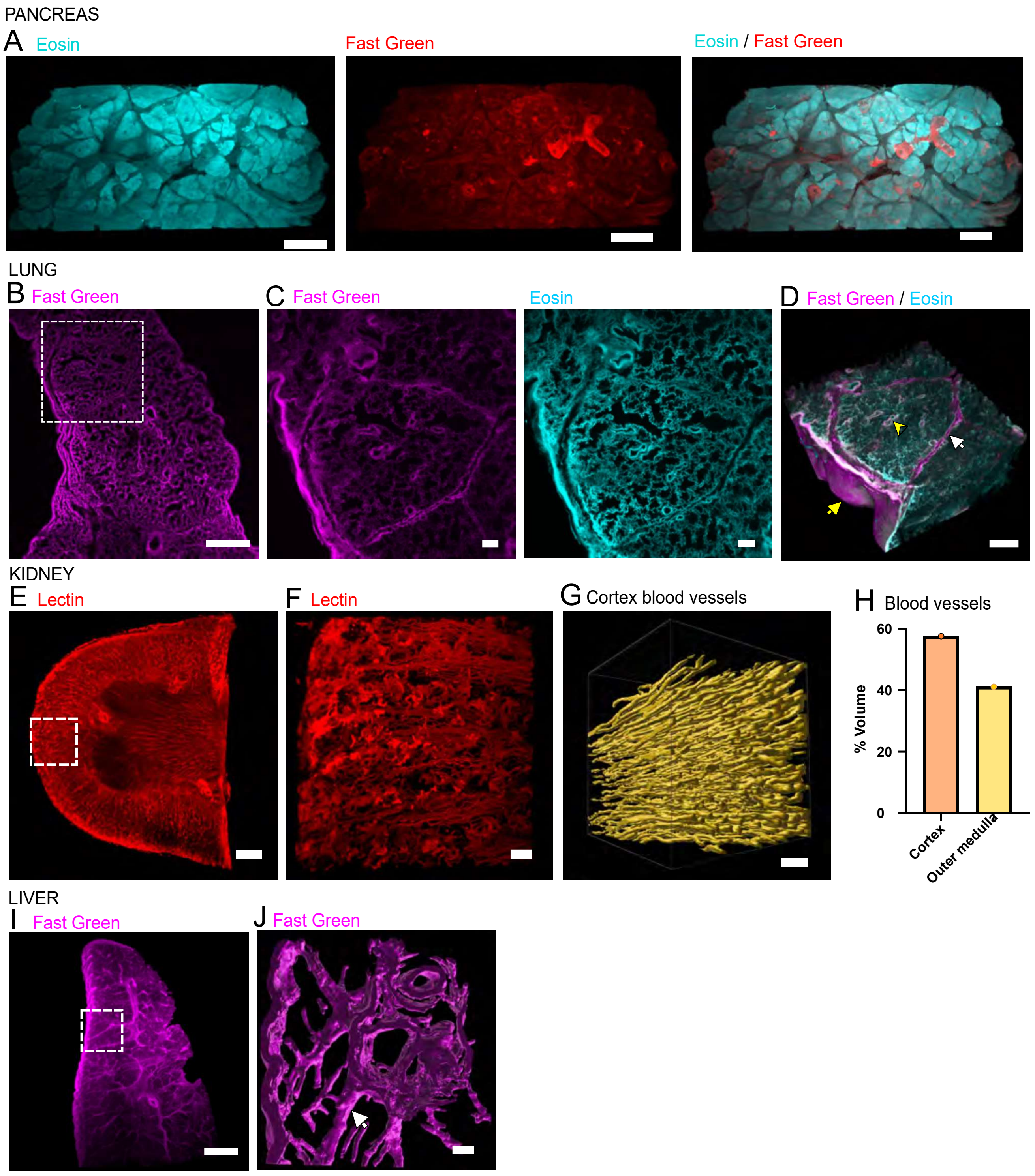
PANCREAS
A Eosin
Fast Green
Eosin / Fast Green
LUNG
B Fast Green
C Fast Green
Eosin
D Fast Green / Eosin
KIDNEY
E Lectin
F Lectin
G Cortex blood vessels
H Blood vessels
% Volume
60
40
20
0
Cortex
Outer medulla
LIVER
I Fast Green
J Fast Green

**Supplementary Figure 3 – 3D quantifications of large organs. (A)** 3D Volume reconstruction of a sample of the pancreas. Pancreatic lobules are labeled with eosin and blood vessels are stained by Fast Green. Scale bar, 2 mm. **(B)** Single plane image from a 3D stack of a lung labeled for collagen with Fast Green. Scale bar, 2 mm. **(C)** Higher resolution acquisition of the inset in panel J labeled with Fast Green and eosin. Scale bars, 500 µm. **(D)** 3D volume reconstruction of a lung lobule enabling the visualization of interlobular septa (white arrow), the pleural surface (yellow arrow) and bronchioles (yellow arrowhead) after Fast Green and eosin staining. Scale bar, 1 mm. **(E)** 3D volume reconstruction of a low magnification image of blood vessels labeled with lectin in a kidney section. Scale bar, 2 mm. **(F)** A higher magnification image of the inset in panel K shows interlobular arteries. Scale bar, 300 µm. **(G)** 3D volume segmentation of interlobular arteries in the renal cortex. Scale bar, 100 µm. **(H)** Volume fraction of blood vessels in the cortex and outer medulla. **(I)** 3D volume reconstruction from collagen labeling in the liver. Scale bar, 2 mm. **(J)** 3D isosurface modeling of the portal triad form collagen staining of the liver, enabling the identification of liver fibrosis (arrow). Scale bar, 300 µm.

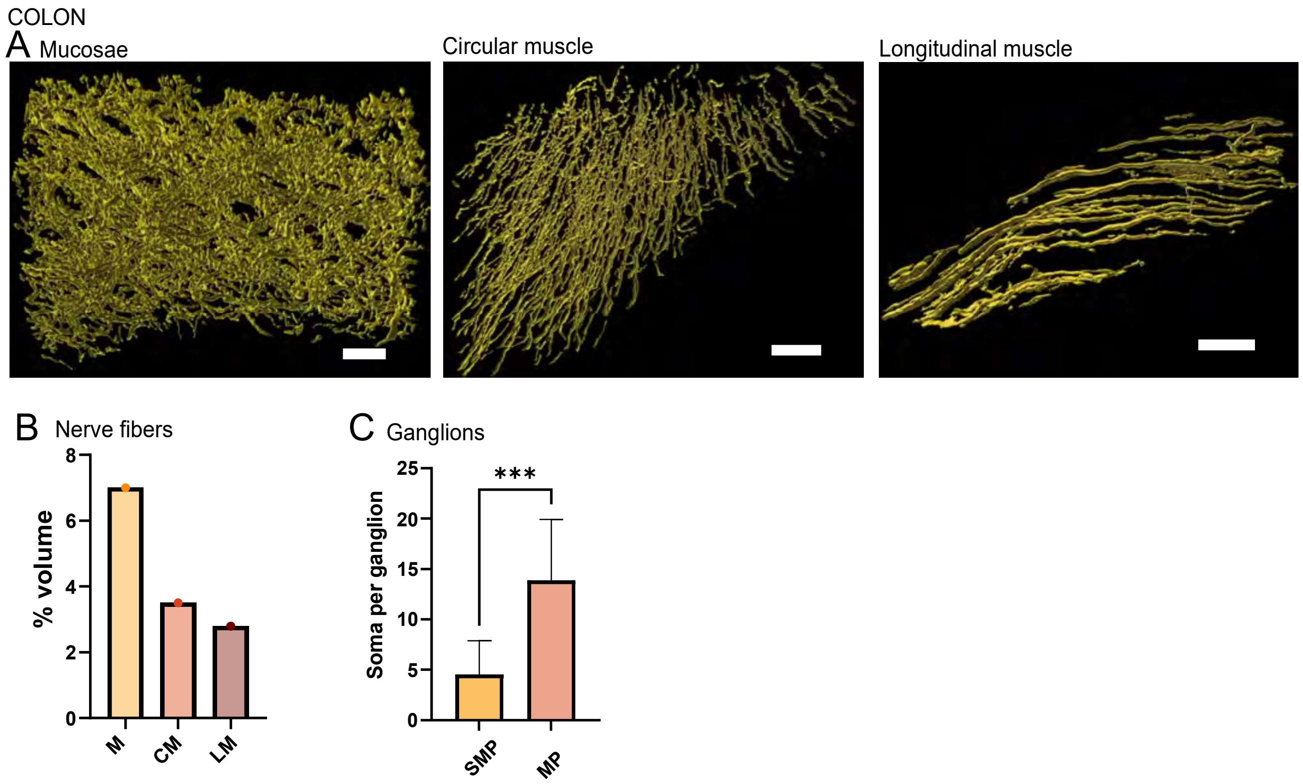


**Supplementary Figure 4 – 3D quantifications of immunolabeling in the colon. (A)** 3D nerve segmentation in the mucosae (M), circular (CM) and longitudinal (LM) muscular layers of the colon. Scale bars, 70 µm, 100 µm and 70 µm respectively. **(B)** Volume fraction of nerve fibers in M, CM, LM. **(C)** Neuronal density quantification a myenteric plexus (MP) and the submucosal plexus (SMP). MP contains a significantly higher number of soma than SP (significant p value<0.05, t-test and mean with standard deviation).

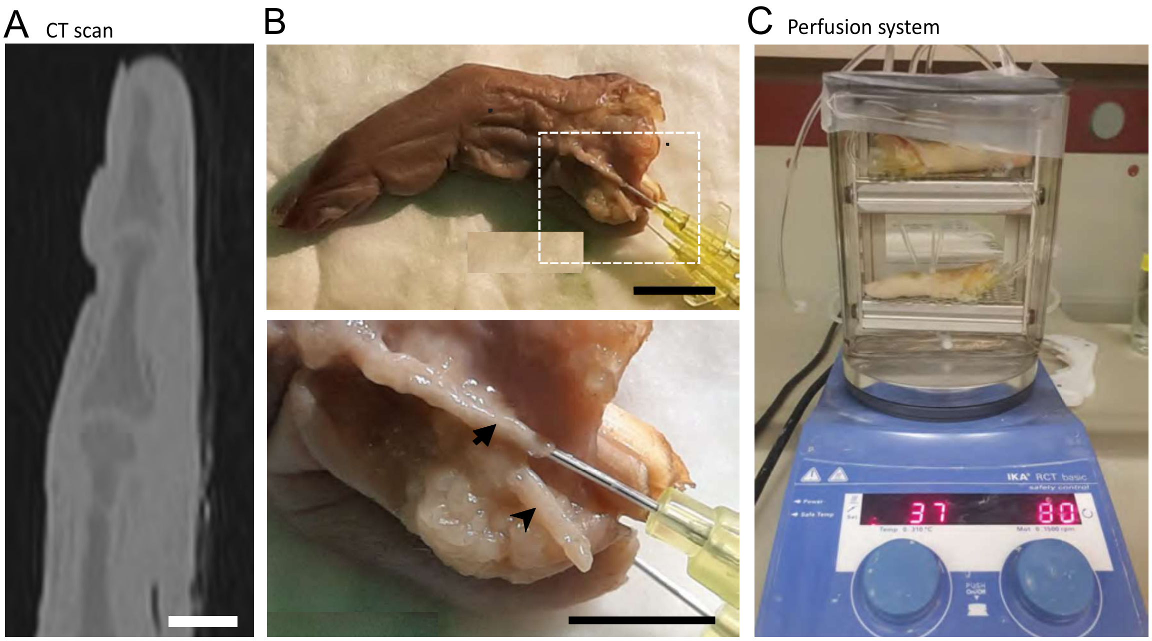


**Supplementary Figure 5 – Processing of human fingers. (A)** Sagittal single plane image of the finger from a CT-scan acquisition after decalcification. Scale bar, 1cm. **(B)** Perfusion was done through a catheter (upper panel) in the ulnar artery (arrow). Lower panel: high magnification inset. The digital nerve is visible next to the artery (arrowhead). Scale bars, 20mm and 10mm respectively. **(C)** Photograph of the closed-loop perfusion setup used to clear the finger.

### Extended data figures legends

**Extended data 1** – Video of a cleared heart related to Fig. 1E.

**Extended data 2** - Video of a cleared kidney related to Fig. 1L.

**Extended data 3 -** Ovary's blood vessels segmentation video related to Fig. 3I. Color scale volume: purple=0.1mm3 to red=0.7 mm3. Scale bar, 1.5 mm.

**Extended data 4** - Video of a 3D volume reconstruction of the lung stained with eosin (cyan) and Fast green (magenta) dyes related to Supplementary Fig. 3D.

**Extended data 5** - Video of a 3D volume reconstruction of the colon related to Fig. 4I. Nerves are labeled for PGP9.5 (red) and blood vessels are labeled for SMA antibodies (green).

**Extended data 6** - Video of a 3D volume reconstruction of the cochlea related to Fig. 5H. Nerves are labeled for beta-III tubulin (cyan).

**Extended data 7** - Video of a 3D volume reconstruction of the finger with SMA immunolabeling (cyan) and Fast Green collagen staining (magenta) related to Fig. 6G.
